# Mass-immigration shapes the antibiotic resistome of wastewater treatment plants

**DOI:** 10.1101/2023.02.27.530348

**Authors:** Lanping Zhang, Bob Adyari, Liyuan Hou, Xiaoyong Yang, Mahmoud Gad, Yuwen Wang, Cong Ma, Qian Sun, Qiang Tang, Yifeng Zhang, Chang-Ping Yu, Anyi Hu

## Abstract

Wastewater treatment plants (WWTPs) are the hotspots for the spread of antibiotic resistance genes (ARGs) into the environment. Nevertheless, a comprehensive assessment of the city-level and short-term daily variations of ARG surveillance is still lacking in WWTPs. Here, 285 ARGs and ten mobile gene elements (MGEs) were monitored in seven WWTPs in Xiamen via high-throughput qPCR (HT-qPCR) for seven days. The average daily load of ARGs to WWTPs was about 1.21 × 10^20^ copies/d, and a total of 1.44 × 10^18^ copies/d was discharged to the environment across the entire city. Interestingly, no daily variations were observed in ARG richness, abundance, and community composition. Stochastic processes were the main force determining the assembly of ARG communities, with their relative importance ranked in the order of influent (INF) > effluent (EFF) > activated sludge (AS). Further analyses indicated that bacteria and ARGs from upstream treatment units played an increasingly dominant role in shaping ARG communities in AS and EFF, respectively, suggesting the importance of mass-immigration of bacteria and ARGs from the source on ARG transport in wastewater treatment units. This emphasizes the need to revise the way we mitigate ARG contamination but focus on the source of ARGs in urban wastewater.

## 1. Introduction

Extensive consumption of antibiotics in humans and livestock exerts a major selective pressure on the environmental microbiome, contributing to the proliferation and spread of antibiotic resistance bacteria (ARB) and antibiotic resistance genes (ARGs) [1]. Therefore, ARB and ARGs have become major global health threats and concerns in environmental systems [2,3]. One Health framework, aiming to understand the mechanism of ARB movement across humans, animals, and the environment, can help guide antibiotic resistance management. Reliable surveillance data lays the foundation for the One Health framework [4]. Previous cross-regional investigations reveal that ARGs are widespread in both highly human-disturbed (e.g., urban city) and human-inaccessible areas (e.g., pristine plateau) [5–9]. Human activities can promote ARG transfer between various environmental interfaces.

Nowadays, WWTPs are widely recognized as the hotspots for ARG proliferation and dissemination [10]. Previous studies showed that the biological treatment processes (e.g., activated sludge) of WWTPs contained a high density of bacteria and various kinds of contaminants (e.g., heavy metals, antibiotics, etc.). These contaminants not only facilitated the proliferation of ARB [11] but also promoted the exchange of ARGs among bacteria [12–18]. Although WWTPs had a relatively high removal efficiency of ARGs for 1–2 log reduction, the effluent still harbored more than 10^7^ copies/mL ARGs [19,20]. The release or recycling of effluents could lead to increasing ARGs abundance and shifting in ARGs community composition in the receiving environments [21–25]. In addition, numerous studies have demonstrated that ARG profiles in WWTP varied at different time scales. For example, at the seasonal scale, the diversity and composition of ARGs were significantly different between summer and winter in eleven WWTPs in China [20]. And the temporal changes of ARG types in effluents were mainly related to the prevalent antibiotics [19]. At the diurnal scale, the variations of ARGs were related to the dynamics of wastewater flow [26,27]. However, the daily variation of ARGs in WWTPs at the city level is still limited. This is important for setting suitable sampling frequencies and monitoring targets to facilitate comparison within and among individual WWTPs. In addition, obtaining this information will help understand the assembly mechanisms underlying ARG spread within a short time, contributing to establishing suitable wastewater-based surveillance approaches in cities.

Stochastic (i.e., dispersal and drift) and deterministic processes (i.e., bio-interaction and environmental selection) collectively contribute to the community assembly [28–30]. However, after decades of debate, the role of stochastic processes in structuring communities has been increasingly appreciated. For instance, the stochastic processes played a central role in structuring ARG communities in urban lakes (e.g., Baiyang Lake and Tai Lake). However, the difference in the relative importance of stochastic processes in Baiyang Lake and Tai Lake was correlated with environmental selection [31]. In addition, Hou and colleagues showed that the stochastic processes played a dominant role in shaping ARG community assembly in urban water bodies under anthropogenic influences [32,33]. And they suggested that the dominant role of stochastic processes may indicate the contribution of HGT to the spread of ARGs in different urban environments. Although many studies explored the influence of various factors (e.g., nutrients, chemical contaminants, operational parameters, etc.) on ARG profiles in WWTPs or bioreactors [17,34–36], the contribution of stochastic processes in ARG community assembly in the different treatment processes of WWTPs is still limited. Understanding the assembly mechanism of microbial communities including ARGs and their hosts is a promising way to provide information for predicting their transport and spread.

Numerous studies reported that the combined effect of physiochemical parameters and biological interaction shapes the ARG profiles [37–39]. Notably, several recent studies have demonstrated that mass immigration via wastewater stream strongly determined the bacterial communities in active sludge [40–42]. However, whether mass immigration (i.e., stochastic processes) plays a role in the ARG community assembly across the wastewater treatment processes is not known. Since influent wastewater flows across different WWTP treatment units, the mass immigration of microorganisms along the flow may shape the microbial and ARG communities in the downstream processes such as the activated sludge and effluents [40].

In this study, we employed HT-qPCR to investigate the daily variation of 285 ARGs and 10 MGEs in INF, AS and EFF of seven WWTPs in Xiamen, China for seven days. We aim to: 1) characterize the daily variations of the abundance, diversity, and composition of ARGs in WWTPs across the whole city; 2) explore the community assembly mechanisms of ARGs and identify the key biotic and abiotic factors shaping the ARG profiles in different wastewater treatment units; and 3) assess the daily ARG mass loads to WWTPs and ARG mass loads in receiving environment at the city level, followed by potential health risk assessment. We assume that stochastic processes will have a higher contribution to the ARG community assembly in INF than in AS and EFF because the latter two will undergo specific environmental pressure (i.e., operation parameters). Among various factors (i.e., biotic and abiotic factors), the immigration of ARGs and microbial communities from upstream (e.g., INF) will affect the ARG profiles in the downstream processes (i.e., AS and EFF). The result of this study will extend our understanding of ARG dynamics as well as the underlying assembly mechanisms in WWTPs, providing fundamental information for the surveillance and control of WWTP ARGs at a city-level perspective.

## 2. Methods and materials

### 2.1 Study area and sample collection

Xiamen is a typical subtropical monsoon city on the southeast coast of China (117°4′E-118°21′E, 24°24′N-24°54′N) with a small range of annual temperature variance (seasonal average temperature 13.7–27.9°C). It covers an area of 1699 km^2^ and has a population of 3.92 million in 2016 [43]. Impressively, the urban land almost doubled to 334 km^2^ and the total amount of municipal wastewater increased from around 210 million tons to 320 million tons during the past 10 years (Xiamen Environmental Protection Bureau, 2007–2016). 92% of the municipal wastewater in Xiamen was treated by seven WWTPs. Among these WWTPs, six WWTPs (listed as W2∼W7) employ the activated sludge technology, while one (i.e., W1) uses a biological aerated filter treatment process. Detailed information on these WWTPs was shown in Table S1. More information about Xiamen city and residents representing the socioeconomic development such as the urbanization rate, per capita disposable income (PCDI), and per capita consumption expenditure was obtained from the Xiamen Bureau of Statistics [44].

The 24-hr composite samples of INF, AS, and EFF were collected from seven WWTPs by using automatic samplers (Hachi, Sigma SD900) for 7 days (from Feb. 28^th^ to March 6^th^, 2016). But there was no sampling event on March 5^th^, 2016 because the staff had the day off. From all WWTPs, six were equipped with one AS tank each and one (W5) was equipped with two tanks. For the first two days, we collected AS samples from the two AS tanks of W5 separately, and one AS composite sample for the subsequent days. Due to the limitations of sampling conditions, a total of 49 INF samples, 49 EFF samples, and 48 AS samples were finally collected. Among these samples, there were 46 pairs of INF-AS-EFF samples. All samples were kept on ice and shipped to the lab within 4 h. For microbial community analysis, 100∼300 mL of INF and 800∼1000 mL of EFF were filtered through the 0.2 μm pore size Sterivex-GP filter (Millipore, Billerica, MA, USA), while AS samples were concentrated by centrifuging at 8,000 rpm for 15 min. All microbial concentrated samples were stored at −80°C before DNA extraction. The concentration of pharmaceuticals and personal care products (PPCPs) was measured following the protocol of EPA 1694 with slight modifications [45]. Here, we classified PPCPs into antibiotics, non-steroidal anti-inflammatory drugs (NSAIDs), stimulants, and other PPCPs, and cited the mass loads of information on each PPCP type from our previous study [45]. All detected PPCPs were listed in **Table S2** and PPCP mass load per capita of each classification were supplied in **Table S3**. The water quality parameters, including chemical oxygen demand (COD), 5-day biological oxygen demand (BOD_5_), total phosphorous (TP), total nitrogen (TN), ammonia nitrogen (NH_4_^+^-N), and suspended solids (SS) of INF and EFF, were provided by the wastewater treatment plants by using national standard methods [46]. Activated sludge performance on specific removal rates of nutrients (TN, TP, and NH_4_^+^-N) and organic matters (indicated as chemical oxygen demand; COD) was calculated as the following equation [47]:

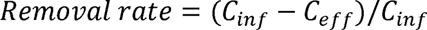

where *C_inf_*and *C_eff_* represent the concentration of each chemical in the INF and EFF (mg/L), respectively.

### 2.2 DNA extraction, 16S rRNA gene amplicon sequencing, and data processing

The genomic DNA of the 146 samples was extracted by using FastDNA SPIN Kit for Soil (Qbiogene-MP Biomedicals, Irvine, CA, USA) according to the manufacturer’s protocol with modifications [43]. The 16S rRNA amplicon sequencing was conducted to characterize the prokaryotic bacterial communities. The 16S rRNA genes in the hypervariable V4-V5 region were amplified with a universal primer set 515F and 907R [48]. The PCR products were then purified and sequenced on Illumina HiSeq 2500 platform (Illumina Inc., San Diego, CA, USA) via a paired-end approach (2 × 250 bp). The raw sequence data can be downloaded from the NCBI short reads archive database under the BioProject number of PRJNA545043.

Raw 16S rRNA paired-end reads were trimmed and assembled using the LotuS pipeline [43,49]. Only high-quality reads that meet the following criteria were retained for subsequent analysis: (1) an average sequence quality > 27; (2) a sequence length > 170 bp; (3) no ambiguous bases; (4) no mismatches to the barcodes or primers; and ([) a homopolymer run < 8 bp. The reads were then checked for chimera and clustered into operational taxonomic units (OTUs) with a threshold of 97% similarity by using UPARSE [50].

### 2.3 HT-qPCR analysis of 285 ARGs and 10 MGEs

A total of 285 ARGs, 10 MGEs, and one 16S rRNA gene were determined on a Warfergen SmartChip real-time PCR system (Wafergen Bio Inc., USA) with the SYBR Green I^®^ chemistry. The targeted ARGs involved the genes that were resistant to all major classes of antibiotics, while those MGEs included 8 transposase and 2 integron genes. HT-qPCR was carried out in a volume of 100 nL reaction system and was under the following thermal cycle condition: initial denaturing at 95°C for 10 min; followed by 40 cycles of denaturation at 95°C for 30 s and annealing at 60°C for 30 s [33,51,52]. The melting curve analyses were automatically generated by SmartChip qPCR software (2.7.0.1). Only the results that meet the following conditions were obtained for statistical analysis: (1) a threshold cycle (Ct) ≤ 31; (2) more than two replicates having been amplified were retained as positive; (3) reaction with one peak; (4) amplification efficiency within 90–110%. The relative copy number of each gene can be calculated using:

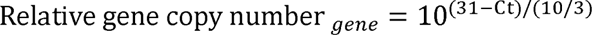

Relative abundance (copies per 16S rRNA gene) and absolute abundance (copies per mL) of ARGs and MGEs was calculated by the following equation:

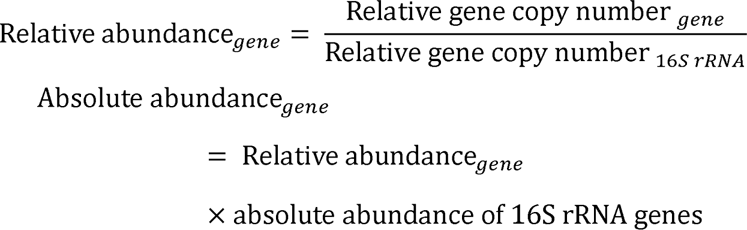

The absolute abundance of the 16S rRNA gene was measured on LightCycler® Roche 480 Real-time PCR instrument (Roche Inc., Basel, Switzerland). A total of 20 μL reaction system consisted of 10 μL LightCycler® 480 SYBR Green I Master Mix (2 ×) (Applied Bio-systems, CA, USA), 0.4 μM of 16S rRNA primer, 1 μg/L bovine serum albumin (Sigma, Steinheim, Germany) and 20 ng of template DNA. The standard curve was processed by 10-fold serially diluted 16S rRNA gene incorporated plasmids with insert length 61 bp bases. The PCR reactions were performed as follows: initial denaturation at 95°C for 3 min, followed by 40 cycles of denaturation at 95°C for 15 s and annealing at 60°C for 1 min and extension at 72°C for 15 s. All qPCR reactions were conducted in triplicate for each sample and negative controls were included.

### 2.4 Daily ARG and MGE mass loads

Based on the total amount of wastewater and the served population, target gene (i.e., ARGs and MGEs) daily mass loads in different WWTPs across the whole city and target gene mass loads per inhabitant are calculated at the city level. ARG or MGE mass loads in different treatment units were following the equations:

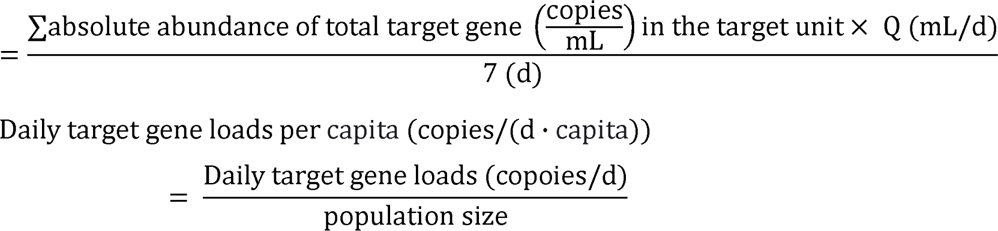

Where Q is the flow rate (mL/d) of the corresponding WWTP. Daily mass loads of ARGs and MGEs per capita in INF and EFF represent the citizen’s daily discharge of ARGs and MGEs to WWTP and potential discharge loads to the environment or release to humans, respectively.

### 2.5 Null model analysis

The null model was utilized to determine the relative contribution of stochastic and deterministic processes to the assembly of ARG and bacterial communities [33]. The principle of the null model can be recognized as follows: the expected similarity of null expected communities was calculated by 1000 random shuffles based on the raw community data. The richness and proportion of species occupancy of the expected community were consistent with the observed community in this step. Then, we calculated the value of the relative importance of the stochastic processes by:

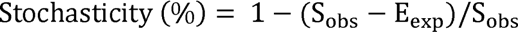

where *S_obs_*means the observed similarity of the original communities, and *E_exp_*means the average similarity of null expected communities.

### 2.6 SourceTracker analysis

The relative contribution of communities (ARGs or OTUs) from different source habitats (i.e., INF, AS, EFF and unknown source) to sink habitats was assessed by using SourceTracker v2.0 with default parameter settings based on previous studies [53–55]. For the sink of AS, the source includes INF and the unknown source, while the sources became INF, AS and the unknown source for the sink of EFF. The unknown source represents the indigenous communities or those from uninvestigated environments. For each sample, the predicted proportion for each of the potential sources was run 5 times to calculate the average proportion. The accuracy of SourceTracker prediction and corresponding standard deviation (SD) within each habitat were evaluated by the consistency between the predicted source and the original habitat. The ARGs and OTUs with an occurrence of less than 1% were discharged during this analysis.

### 2.7 Statistical analysis

Daily variations of ARG richness and abundance for seven days were assessed by the coefficient of variance (CV) and were calculated as the following formula: CV = SD/the mean value of ARG richness or abundance. The time-dependent ARG and bacterial community structures were analyzed by using a time decay model. The time decay model takes the time interval matrices between samples as the independent variable, and the β-diversity matrices based on the Bray-Curtis distance between samples as the dependent variable. The variables were fitted with a linear model and the obtained slope and R^2^ were used to evaluate the collinear relationship.

The β-diversity patterns of ARG and bacterial communities were revealed by nonmetric multidimensional scaling (NMDS) analysis based on Bray-Curtis distance matrices. The significance of the differences within ARG and bacterial communities among wastewater treatment units or WWTPs were performed by permutational multivariate analysis of variance (Adonis) and analysis of similarity (ANOSIM).

VPA was used to distinguish the relative effects of biotic, abiotic factors, and MGEs on the variance of β-diversity of ARG communities in each treatment process [48]. The biotic factors included the first two NMDS axes of the bacterial communities and the corresponding source proportions of ARG and bacterial communities predicted by SourceTracker. Here, we only included the source proportions in the analysis of ARG communities from AS and EFF. The abiotic factors included physicochemical parameters of INF and EFF and the nutrient removal rate (i.e., TN, TP, NH_4_^+^-N, and COD) of WWTPs. Here, MGEs were included as an independent factor because they were commonly recognized as an important driver causing the HGT of ARGs. Prior to VPA, the multicollinearity between variables in each factor group was assessed by the variation inflation factor (VIF), using “vifstep” function from the R package *usdm* [56]. The variables with VIF < 10 were retained. To quantify the relative importance of an individual element of factors, HP analysis was performed via the “rdacca.hp” function from the R package *rdacca.hp* [57].

Statistical analysis and visualization were conducted using R 4.1.0 with the R package *phyloseq* [58], *vegan* [59], and *ggplot2* [60].

## 3. Results

### 3.1 A full profile of ARGs in WWTPs across the whole city

A total of 208 ARG subtypes were detected in INF, AS, and EFF of seven WWTPs of Xiamen, corresponding to 8 major ARG types (i.e., aminoglycoside, beta-lactam, chloramphenicol, MLSB, multidrug, sulfonamide, tetracycline, and vancomycin resistance genes). The number of ARG subtypes in different treatment units decreased as follows: INF (94∼141) > EFF (31∼115) > AS (44∼98) (**Fig. S1A**). 156 of 208 ARG subtypes (75%) were shared by three habitats (i.e., INF, EFF, and AS) (**Fig. S1B**). The shared ARGs mainly belonged to beta-lactam (43 ARG subtypes), multidrug (33), aminoglycoside (31), and MLSB (23) resistance genes.

The results indicated that the relative abundance of ARGs in INF was the highest (average 0.56 ± 0.22 copies/16S rRNA gene), and then decreased more than 50% in AS (0.20 ± 0.09 copies/16S rRNA gene). But in EFF this value increased again to 0.33 ± 0.16 copies/16S rRNA gene (**Fig. 1A**). Being inconsistent with this trend, AS had the highest absolute abundance of ARGs (average 4.14 × 10^9^ copies/mL), followed by INF (1.34 × 10^8^ copies/mL) and then EFF (2.40 × 10^6^ copies/mL) (**Fig. 1B**). According to the absolute abundance of ARGs, aminoglycoside (ranging from 2.70 × 10^4^ to 5.36 × 10^9^ copies/mL), multidrug (3.49 × 10^4^∼4.82 × 10^9^ copies/mL), MLSB (5.81 × 10^3^∼9.03 × 10^8^ copies/mL) and beta-lactam (9.90 × 10^3^∼3.06 × 10^9^ copies/mL) resistance genes were the top 4 dominant ARG types.

**Figure 1.**
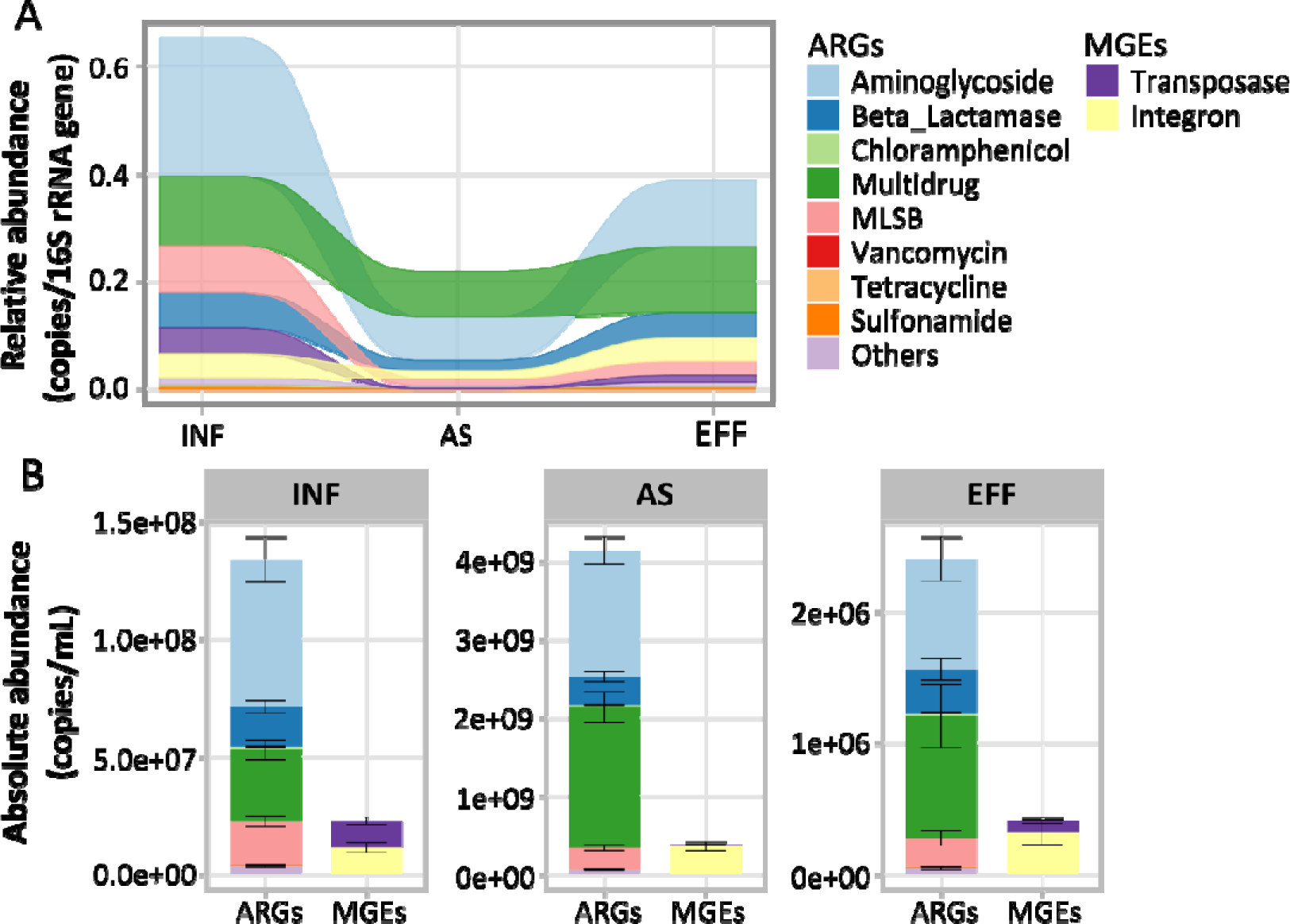
Relative abundance (A) and absolute abundance (B) of ARGs and MGEs in different wastewater treatment units. MLSB stands for Macrolide-Lincosamide-Streptogramin B.

As for MGEs, the relative abundance in INF was the highest value (average 0.092 ± 0.05 copies/16S), followed by EFF (0.055 ± 0.07copies/16S) and AS (0.016 ± 0.01 copies/16S) (**Fig. 1A**), while the absolute abundance is in the following order: AS (average 3.81 × 10^8^ copies/mL) > INF (2.26 × 10^7^ copies/mL) > EFF (4.10 × 10^5^ copies/mL) (**Fig. 1B**). Notably, among MGE subtypes, the relative abundance of integrons in EFF increased to a level comparable to that in INF. Spearman correlation analysis indicated that the relative abundances of MGEs and ARGs were significantly correlated with each other in both INF and EFF (r ≥ 0.50, *P* < 0.001), but not in AS (**Fig. S2**). In addition, the relative abundance of transposases correlated significantly with the majority of ARG types in both INF and EFF (*P* < 0.05). Interestingly, the relationship between integrons and ARG types was stronger in INF than in EFF (**Fig. S3**).

### 3.2 Daily variations of ARG abundance, diversity, and composition in different treatment units

NMDS ordination analysis indicated that both ARGs and bacterial communities tended to cluster based on the sample habitats (i.e., treatment units) (**Fig. 2A & C**). The treatment unit (R^2^ = 0.21, *P* < 0.001) and WWTPs (R^2^ = 0.25, *P* < 0.001) showed similar effects on bacterial community compositions of WWTPs based on Adonis tests. However, the overall variance of the ARG communities driven by the treatment unit (R^2^ = 0.35, *P* < 0.001) was much higher than that of WWTPs (R^2^ = 0.14, *P* < 0.001). Significance differences have been observed in both ARG and bacterial communities between EFF and INF (ARGs: R^2^ = 0.19, *P* < 0.01; bacteria: R^2^ = 0.14, *P* < 0.01), EFF and AS (ARGs: R^2^ = 0.20, *P* < 0.01; bacteria: R^2^ = 0.11, *P* < 0.01), as well as INF and AS (ARGs: R^2^ = 0.43, *P* < 0.01; bacteria: R^2^ = 0.26, *P* < 0.01). However, the differences in both ARG and bacterial communities between EFF and INF, and EFF and AS were much smaller than those between INF and AS. This indicated the significant shifts of ARG and bacterial communities from INF to AS. In addition, bacteria and ARG communities showed pronounced spatial differences across the treatment units, especially in AS (ARGs: R^2^ = 0.66, *P* < 0.001; bacteria: R^2^ = 0.78, *P* < 0.001) (**Fig. S4**). However, Procrustes analysis revealed a similar and significant β-diversity pattern between ARG and bacterial communities within each treatment process (**Fig. S5**).

**Figure 2.**
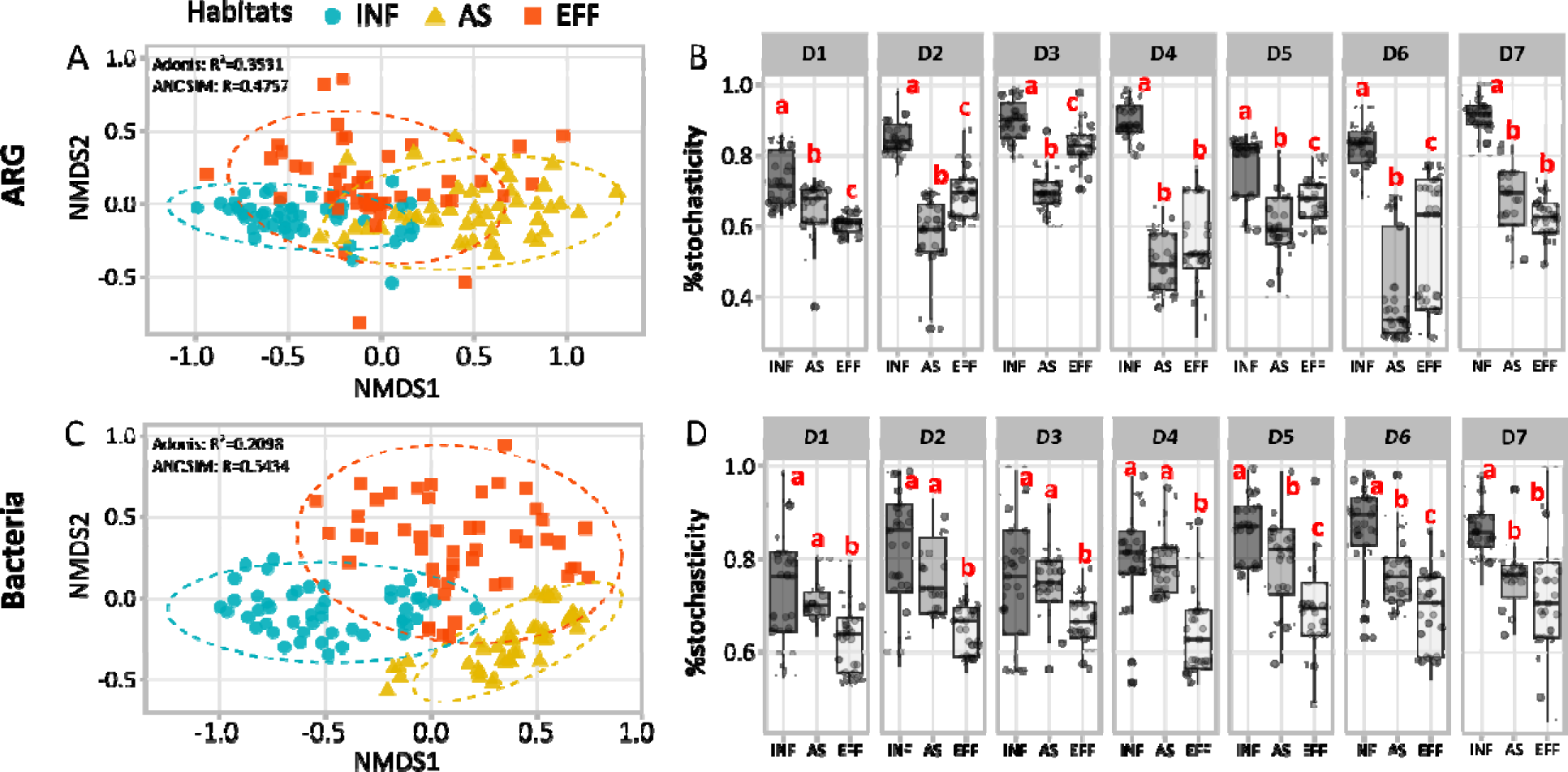
Non-metric multidimensional scaling (NMDS) ordination based on Bray-Curtis distance showing the distribution pattern of ARG communities (A) and bacterial communities (C) in all WWTPs across the entire city. The ellipses were drawn at the 95% confidence level. Abundance-based β-null model was applied to assess the relative importance of the stochasticity process in governing the assembly of ARG (B) and bacterial communities (D) in each treatment unit during the sampling period (from day 1 to day 7). Significant differences were indicated by different letters among different units of the treatment process.

The daily variation of the ARG communities among different treatment units during the sampling period was presented in **Fig. 3**. The daily variation of ARG richness showed statistically significant differences between INF and AS, and INF and EFF (Wilcoxon test; *P* < 0.05). The relative abundance of ARGs exhibited much smaller daily fluctuations compared to the absolute abundance based on the CV analysis (**Fig. 3A**). Among different treatment units, AS had the greatest variation in ARG richness over time but the smallest variation in ARG relative abundance and OTU richness. Based on the time decay model, we found how the dissimilarity of ARG community composition changes with time. No linear association between the dissimilarity of ARG community composition and time has been observed. However, the dissimilarity of the bacterial community composition showed a strong positive correlation with time (**Fig. 3B**). This indicated that the daily variation of the ARG community within the same treatment unit was stable.

**Figure 3.**
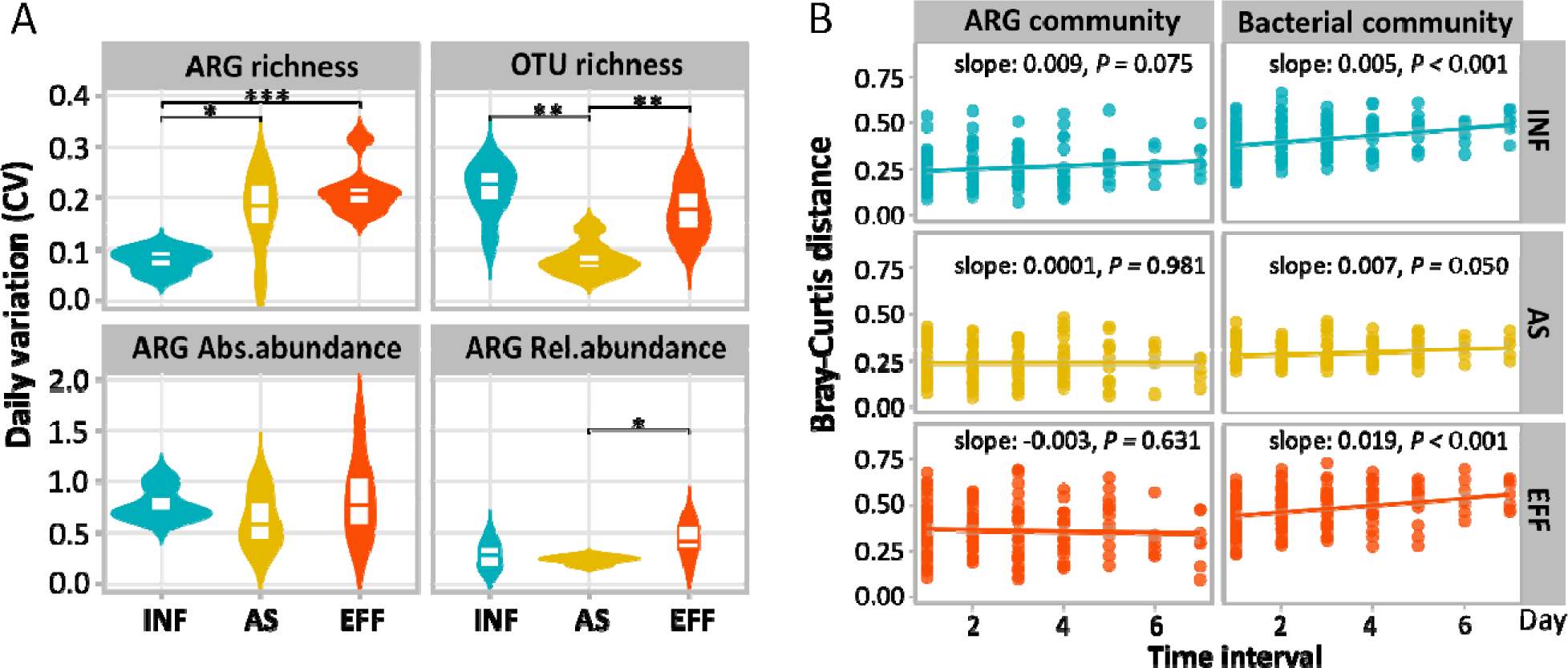
Daily variation of ARGs and bacteria at different wastewater treatment units. (A) Coefficient of variation (CV) of richness and abundance of overall ARG community and OTU richness, (B) time-decay relationships of the compositional dissimilarity of ARG and bacterial communities at different time intervals. Significant differences among different units of the wastewater treatment process were assessed by paired Wilcoxon test and indicated as follows: \*\*\**P* < 0.001, \*\**P* < 0.01 and \**P* < 0.05. Lines represent the fitted linear regression curve. The shaded areas represent the 95% confidential interval of the fitted linear regression curve.

### 3.3 Immigration of ARGs across different treatment units

SourceTracker analysis conducted the prediction of the source habitats (i.e., the treatment units) for ARG communities in 78% (108/138) samples and for bacterial communities in 91% (126/138) samples successfully (**Figs. S6A and S7A**). Specifically, as for ARG communities, 89% samples (41/46), 89% samples (41/46), and 57% samples (26/46) were predicted to be from INF, AS, and EFF with a probability of 73%±15%, 79%±14%, and 62%±16%, respectively. This supported the findings of NMDS ordination that the samples tended to cluster by their habitats (**Fig. 2A & C**), suggesting that each habitat had unique characteristics of ARG communities. However, as the wastewater flows between different treatment units, the immigration process of the upstream communities (i.e., INF) contributed to downstream communities (e.g., AS and EFF). For instance, the contribution of INF to the ARG community downstream (i.e., AS and EFF) increased greatly along with treatment units from 25.5% in AS to 49.6% in EFF. AS also contributed to 24.3% of the ARG community in EFF (**Fig. S6B**). As a comparison, the contribution of INF to the bacterial community in AS was 56.1%, while INF and AS had similar contributions (∼33%) to the bacterial community in EFF (**Fig. S7B**).

### 3.4 Assembly mechanisms and controlling factors underlying ARGs in WWTPs

The null model analysis showed that stochastic processes played a dominant role in shaping the ARG communities in WWTPs (the average stochastic ratio > 50%), and the relative importance of stochastic processes ranked as follows: INF (average 83.7% ± 9.3%) > EFF (64.6% ± 13.9%) > AS (58.8% ± 13.9%) (**Fig. 2B**).

VPA results showed that biotic (i.e., local bacterial communities and immigration of bacteria and ARGs), abiotic (i.e., physicochemical parameters) factors and MGEs could explain 36.5%∼40.5% of the total variance of ARG communities in different treatment units (**Fig. 4A, C, & E**). Although the pure effects of abiotic factors and MGEs are relatively low in three treatment units, the interaction effect of these factors was high in INF. Notably, the pure effects of biotic factors on the total variance of ARG communities increased greatly as wastewater flowed to the downstream treatment units (5.5% in INF, 13.2% AS and 23.5% in EFF). The relative importance of an individual factor to the variance of ARG communities was assessed by HP analysis (**Fig. 4B, D, & F**). The results indicated that bacterial immigration from INF is the most important to the total variance of the ARG communities in AS. And ARG immigrations from INF and AS were the top two important factors affecting ARG communities in EFF. Besides, the pure effects of MGEs on ARG communities decreased along with treatment units.

**Figure 4.**
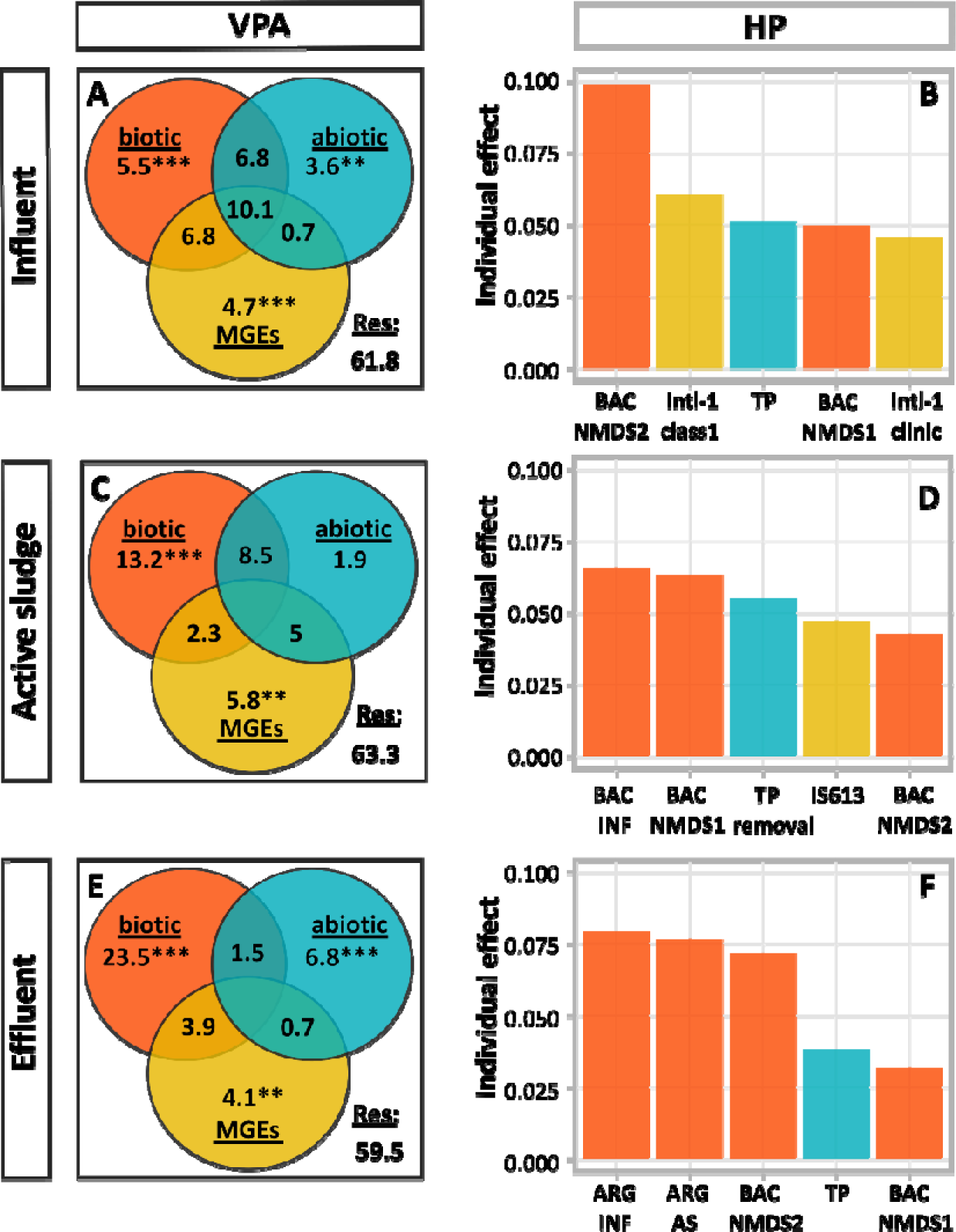
Variation partitioning analysis (VPA) of β-diversity variation of ARG communities among biotic, abiotic, and MGEs groups as well as their interactions (A, C, E). The absence of any values for the three interactions in Figure C & E implies that the interaction effect was 0. MGEs are considered to be a unique class of factors that contribute to the assembly of the ARG community, and therefore are separated from the biotic group as a single factor. The top five variables with the highest variance explained to β-diversity variation of ARG communities in each treatment unit are indicated by hierarchical partitioning (HP) analysis (B, D, F). The abbreviation of BAC NMDS1 and BAC NMDS2 represent the first two axes of NMDS results of the bacterial community in each treatment unit, and the abbreviation of BAC INF, BAC AS, ARG INF, and ARG AS represents the source of the bacterial community or ARG community (upstream treatment units such as INF or AS). Significant differences were indicated as follows: \*\*\**P* < 0.001, \*\**P* < 0.01 and \**P* < 0.05.

### 3.5 Daily mass loads of ARGs to the urban environment at the city level

Daily mass loads received by WWTPs across the whole city were 1.21 × 10^20^ copies/d ARGs and 2.01 × 10^19^ copies/d MGEs. And WWTPs discharged 1.44 × 10^18^ copies/d ARGs and 2.43 × 10^17^ copies/d MGEs to the receiving environment. The citizen’s daily ARG and MGE mass loads to WWTPs can be calculated accordingly, which are up to 3.13 × 10^13^ copies/d/capita and 5.20 × 10^12^ copies/d/capita, respectively. And potential discharge loads to the environment or releasing to humans from WWTPs were 3.72 × 10^11^ copies/d/capita, 6.31 × 10^10^ copies/d/capita, respectively (**Fig. 5A & B**).

**Figure 5.**
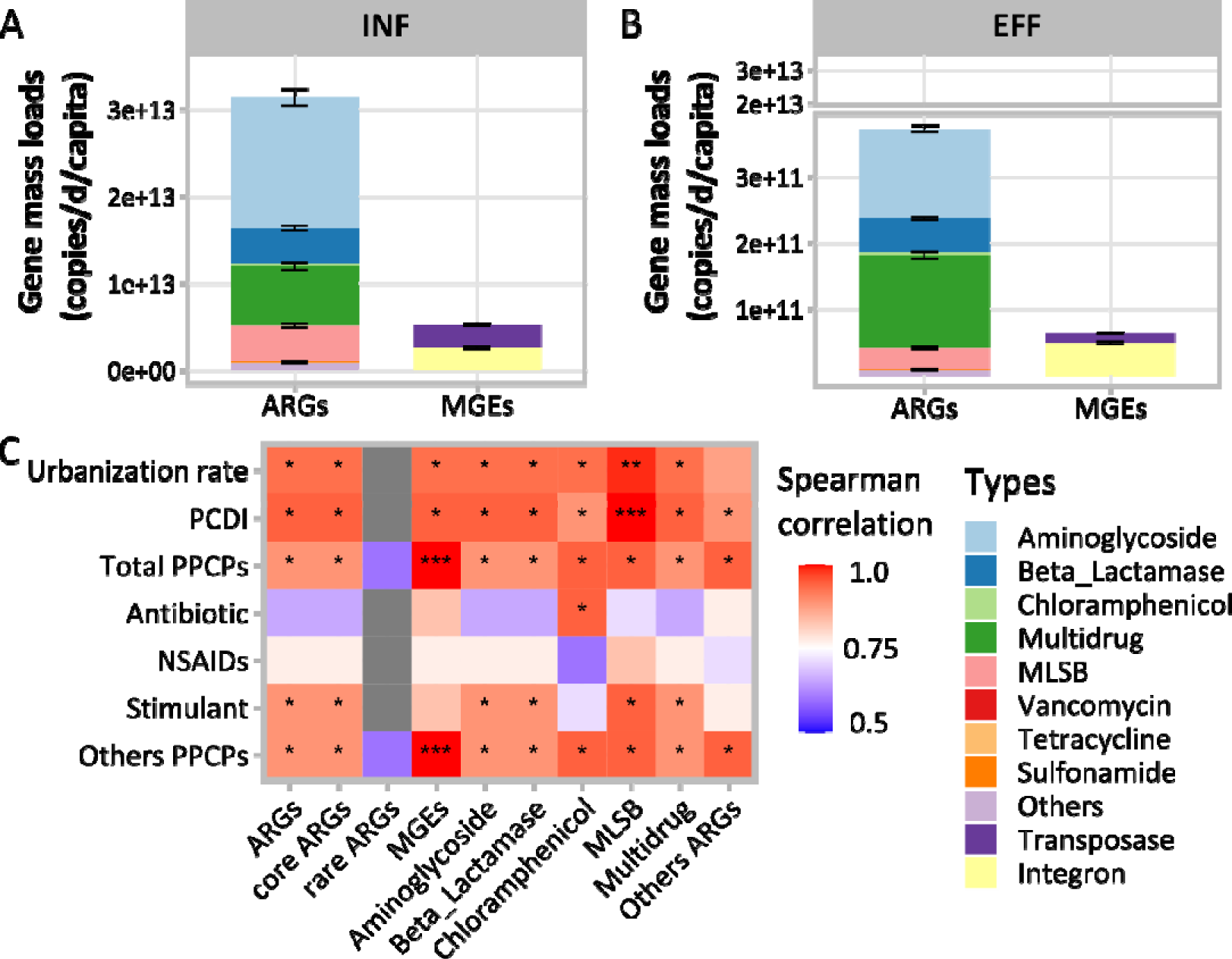
The daily ARG and MGE receiving loads per capita by WWTPs (A) and discharge loads from WWTPs (B) in Xiamen, China. Spearman correlation analysis was made between major types of the daily discharge ARG loads per capita and the daily PPCP loads in WWTPs and social development factors of the city (C). Significant differences were indicated as follows: \*\*\**P* < 0.001, \*\**P* < 0.01 and \**P* < 0.05.

Spearman correlation analysis indicated that total PPCPs mass loads, other PPCPs mass loads, and socioeconomic factors (i.e., urbanization rate and PCDI) were significantly correlated with six major types of ARG mass loads and the MGE mass loads, while antibiotics and NSAIDs had no significant correlation with any types of ARGs (**Fig. 5C**). In addition, MGE loads had the highest correlation coefficient with the total PPCPs mass loads.

## 4. Discussion

To facilitate the monitoring and control of ARG dissemination, setting up a wastewater-based surveillance approach as well as understanding the ecological processes and driving forces of the ARG communities along the wastewater treatment processes are required. Unlike the previous studies that mainly focused on the fate of ARGs in WWTPs, we targeted the variation and transport of ARGs in different wastewater treatment units. Here, we found the stable daily variance of ARGs at the city level through large-scale and continuous daily sampling. Furthermore, stochastic processes were the dominant assembly mechanism of ARG communities in WWTPs, although the importance in each treatment unit varied. Among the driving factors, the immigration of bacteria and ARG in the upstream treatment unit significantly shaped the downstream ARG community. At last, we estimated the daily mass loads of ARGs to the WWTPs and the ones released from WWTPs at the city level.

### 4.1 Stable daily ARG profile and loads throughout WWTPs inspired ARG surveillance strategies

WWTPs are recognized as the breeding reservoirs for ARG dissemination, which harbor ARGs with high diversity and abundance. However, there is limited information regarding the continuous ARG pollution at various WWTP units during wastewater treatment operations. A comprehensive understanding of the daily dynamics and profile of ARGs throughout WWTP units is highly needed to build wastewater-based surveillance approaches for One Health. Here, we found that the composition of the daily ARG communities tended to be stable throughout seven days from all WWTPs in the same city (**Fig. 3B**). Interestingly, the ARG richness and total ARG relative abundance in INF showed low daily variations (**Fig. 3A**). This indicated that the ARGs profiles introduced by sewage formed a relatively stable pattern across entire city, compared to the EFF, which is affected by daily treatment process variability [61]. Another reason for such abundant and stable ARGs is the plateau effect of microbes. That is, microbes have reached the highest carrying capacity of ARGs in special environment. The plateau effect was frequently observed in field irrigation experiments [24,62]. For instance, the relative abundance of *sul1* in fields irrigated with untreated wastewater for 1.5 years was similar to that in fields irrigated with wastewater for 10 years [62]. This phenomenon requires continuous stimulation of high ARGs to stabilize [24]. Furthermore, the high stability of ARG profiles in INF indicates the locational reproducibility, suggesting that applying a low sampling frequency of sewage would be useful for ARG surveillance in global WWTPs. Similar phenomenon, such as shifts and increases in variability of ARGs from INF to EFF, have been observed in other studies [61,63]. The relatively higher daily variation of ARGs in EFF also presents advantage in assessing WWTP performance and providing a measure of loads and potential associated exposures to the receiving environment. Additionally, it may be better than by INF surveillance alone for detecting and dealing with anomalies of concern [61]. Another landmark study in Hong Kong WWTPs with similar climate and geographical location showed no distinct seasonal variations over 9 years span. However, the ARG composition in AS shifted every 2∼3 years [64]. In this study, no such turnover period was captured, and it is unclear to what extent the observed patterns are generalizable across all WWTPs. Future research would focus long period sampling for explore the ARG turnover pattern and dynamics across each treatment processes.

Our study revealed that the relative abundance of ARGs showed higher dynamic robustness than the absolute abundances of ARGs, which is consistent with a previous study [27]. This study suggested that applying the ARG relative abundance as the index can avoid minor variations in sample processing. In addition, our hourly composite sampling over a 24-hour period reduced the impact of accidental residential water use and provided a closer representation of daily discharge of ARGs into WWTPs [26,27]. Overall, the above results provided novel insights into sampling and monitoring strategies for antibiotic resistance surveillance, particularly for monitoring anomalous genes of clinical concern in WWTPs, that is, short-term sampling for ARGs profiles can provide representative information for the distribution of ARGs in WWTPs (at least a week).

### 4.2 Immigration process determined the assembly of the ARG communities

The ecological mechanisms of ARG community assembly have been investigated mainly in urban semi-natural environments, such as urban ponds and reservoirs [33,37,38,65]. However, little is known about the ARG community assembly processes in engineered environments. Here, the central role of stochastic processes in the ecological assembly processes of ARG communities in WWTPs was revealed. It is consistent with our previous study with a small sample size [33]. The prevalence of such high stochasticity in ARG communities throughout WWTPs in the current study may be attributed to other factors beyond the effects of small-scale habitats and homogeneous physicochemical conditions [65,66]. The high correlation between the abundance of ARGs and MGEs in INF suggested that HGT may play an important role in the spread of ARGs (**Fig. S3**). HP analysis found that MGEs were the second most important factor in determining the ARG communities in INF (**Fig. 4B**). Indeed, large amounts of nutrients provided beneficial conditions for nourishing microbes as well as pathogens, which could facilitate cell contact and their HGT of ARGs there [12,67].

In AS, we found that the immigration of bacteria from INF and the local bacterial communities was more vital compared to the other two factors (i.e., MGEs and abiotic factors) (**Fig. 4C & D**). In a previous study, the bacterial community was considered to be the key factor that drove the ARG profiles in the high anthropogenic environment [51]. The variation in the microbial community composition was concurrent with stable metabolic functional structure and performance [68]. In the AS bioreactor, to maintain high efficiency in nutrients removal, the growth of species with desired function or high adaptability is facilitated via operation parameters such as solids retention time and recycle ratio, which is a deterministic process that may affect the proliferation of ARGs [36,40,69]. This indicated the importance of VGT for ARG reproduction in AS. Recently, researchers confirmed that the VGT is significantly involved in the ARG community of AS via single-cell microfluidics [70,71]. Thus, the existing understanding suggested deterministic factors, such as the design and operation of AS bioreactors, are suggested to contribute to the AS microbial communities. However, we showed that microbial immigration from upstream (stochastic processes) mainly controls the ARG communities in AS. Similarly, Dottorini et al. found that the influent microbial community immigration determined the assembly of the AS microbial communities [40]. Previously, INF microbial communities were often considered as the seeding for the AS communities. However, here, mass immigration is responsible for both the presence and abundance of ARGs in the AS, especially for those species with ARGs that are not fit for the function of AS.

In EFF, the low bacterial density and diversity, as well as the low concentration of nutrients and antibiotics limited the frequency of VGT occurrence, while the high potential for natural transformation of ARG via HGT was confirmed [72,73]. Interestingly, our analysis revealed that the immigration of upstream ARGs had the highest effects on shaping the ARG communities, followed by the upstream and local bacterial communities and then MGEs (**Fig. 4F**). Tong et al. think bacteria host with ARGs may survive better compared with the bacteria without ARGs during the treatment processes [18]. This explained why the immigration of ARGs played a more important role in controlling the ARG community than the bacterial community. This also tells us the source ARG community is essential for the ARG discharges from WWTPs to the environment, challenging us to revise our understanding to control ARGs in sewer systems before WWTPs. Besides, the contribution of the unknown source for the ARG community in AS and EFF decreased along with the treatment units, indicating the limitation of the general source library of ARGs in WWTPs. Future studies focusing on providing more source information into the analysis and therefore increasing the range of identifiable sources in WWTPs are needed.

Although researchers found that EFF largely reflected AS and INF by microbes escaping settling, it is hard to quantify the influence from upstream [61]. We reported the first result highlighting the importance of the immigration of bacteria and ARGs from the source/upstream in shaping the downstream ARGs communities in WWTPs. Our results revealed that the shift of the ARG community structure is mainly affected by the immigration from upstream treatment units, but the immigration of the bacterial and ARG communities contributed differently. WWTPs were continuously fed by upstream sewage sources and were open ecosystems [40,74]. The INF including bacteria was considered to be the main source of biomass in AS and may influence the diversity, abundance, and assembly of local microbial communities microbes via microbial immigration [40–42,75,76]. Although microbial immigration constantly contributes to community assemblage in natural and engineered microbial ecosystems, the impact of microbial immigration varied due to different immigration intensities [77,78]. ARGs are prone to persist in the receiving environment once introduced [24]. For example, effluents of WWTPs rapidly stabilized the antibiotic resistome in receiving freshwater, even if bacterial community composition was not yet stable, implying the occurrence of ARGs and pathogens downstream was affected by the exchange of ARGs from EFF [22,77,79]. However, no evidence was found of taxa that could drive the stabilization of ARGs in receiving environment [77]. Recently, coupling a mass balance model with qPCR allows the calculation of the effects of treatment performance and hydrologic characteristics on ARG communities in receiving environment [21,23]. However, the relative importance of the immigration of bacterial and ARG communities in shaping ARG communities remains veiled in engineered ecosystems. Here, we found that the immigration of bacteria and ARG community determined the ARG communities in AS and EFF, respectively. The results fill up the gap by providing basic information on the dissemination of ARGs throughout WWTP treatment units and novel insights for policy maker to consider the role of mass immigration when developing strategies to control ARGs.

### 4.3 WWTPs as ARG dissemination source under socioeconomic development at the city level

With the rapid development of industry, antibiotics are widely used in animal husbandry and disease treatment, thus making the resistance gradually become a serious problem. In our study, we found that socioeconomic factors are closely related to ARG pollution across the whole city from the perspective of ARG loads. Population size is the key factor driving fluoroquinolones, macrolide antibiotics, sulfamethoxazole, chloramphenicol, and daily loads of the corresponding ARGs in INF [80]. Xiamen is a coastal city with rapid industrialization and economic development. Two WWTPs with the highest ratio of domestic wastewater in INF contained the highest relative abundance of total ARGs. This indicated that domestic wastewater which is closely associated with population size is the main source of ARGs. Our previous study found that socioeconomic factors (i.e., urbanization rates) are significantly correlated with PPCP daily mass load per capita [45]. Here, a positive correlation between two socioeconomic factors (i.e., urbanization rate and PCDI) and daily ARGs per capita was verified. This suggested that higher human activity facilitates ARG pollution in highly developed area.

A study found that two municipal WWTPs in South Korea covering < 40% of the population in Xiamen received average ARG daily loads of around 10^20^ copies/d and discharged 10^16^∼10^18^ copies/d to the environment [81]. The total ARG daily mass loads in Xiamen WWTPs in China were in the same range, suggesting the burden of ARG pollution in highly developed cities. ARG and MGE mass loads are another vital indicators for ARG dissemination assessment [24]. Thus, despite the fact that two orders of magnitude of ARGs were removed during the treatment processes of WWTPs, a massive amount of ARGs and MGEs were released into the environment across the whole city, leading to the subsequent wide-spread and high risk of exposure of ARGs to humans.

## 5. Conclusion

After characterizing the ARG dynamics throughout the wastewater treatment units in all WWTPs in a city for a week, the following conclusions were therefore drawn:

1. Short-term sampling for ARGs profiles can be representative enough for the distribution of ARGs in WWTPs in the longer term (at least a week), such as ARG richness, relative abundance, and absolute abundance of total ARGs, as well as the compositional dissimilarities of the ARG communities in a week showed little daily variation, especially in INF.

2. Stochastic processes in assembly ARG communities were dominant in three treatment units, and the relative importance of stochastic processes in different treatment units ranked as follows: INF > EFF > AS. The immigration of the bacterial communities upstream and the local bacterial communities played an important role in shaping the ARG communities in AS. The immigration of ARG communities from upstream in WWTPs combined with the local bacterial communities contributed to the ARG dissemination in EFF. This emphasizes the need to revise the way we mitigate ARG contamination but focus on the source in the urban environment.

3. We assessed the total ARG daily mass load received by and discharged from WWTPs at the city level, filling the knowledge gaps on the total amount of ARGs in the urban water cycle. We have linked the socioeconomic factors at the city level with the ARG pollution from WWTPs, providing an assessment of human exposure to ARGs for highly developed cities.

Overall, this study highlighted the daily pattern of ARGs throughout the wastewater treatment units and the importance of mass immigration of ARGs and bacteria from upstream on the formation of ARG communities downstream. This provides the fundamental information for establishing wastewater-based surveillance strategies and ARG ecological behavior in WWTPs.

## Author contributions

**Lanping Zhang**: Conceptualization, Investigation, Methodology, Data Analysis, Software, Visualization, Writing - original draft. **Bob Adyari**: Validation, Software, Writing - review & editing. **Liyuan Hou**: Validation, Writing - review & editing. **Xiaoyong Yang**: Investigation, Writing - review & editing. **Mahmoud Gad**: Writing - review & editing. **Yuwen Wang**: Investigation, Methodology. **Cong Ma**: Investigation, Resources. **Qian Sun**: Methodology, Resources. **Qiang Tang**: Writing - review & editing, Funding acquisition. **Yifeng Zhang**: Writing - review & editing. **Chang-Ping Yu**: Resources, Funding acquisition. **Anyi Hu**: Conceptualization, Data curation, Validation, Writing - review & editing, Supervision, Project administration, Funding acquisition.

## Acknowledgement

We thank Dr. Fuyi Huang and Miss. Xinyuan Zhou for their assistance with the methodology. This work was supported by the National Key R&D Program of China (2022YFE0120300), the National Natural Science Foundation of China (U1805244 and 31870475) and CAS Key Laboratory of Urban Pollutant Conversion Joint Research Fund (KLUPC-2021-2). M. Gad was supported by the Science, Technology & Innovation Funding Authority, Egypt (44201).

