## supplemental files for "Mass-immigration shapes the antibiotic resistome of wastewater treatment plants"

**Table**

**Table S1**: The process composition of seven wastewater treatment plants.

| **WWTPs** | **Process composition** |
| --- | --- |
| W1 | Primary sedimentation, biological aeration tank, UV disinfection |
| W2 | Primary sedimentation, oxidation ditch, secondary sedimentation, UV disinfection |
| W3 | Primary sedimentation, oxidation ditch, secondary sedimentation, UV disinfection |
| W4 | Primary sedimentation, A^2^/O, secondary sedimentation, chemical disinfection |
| W5 | Primary sedimentation, A^2^/O, secondary sedimentation, UV disinfection |
| W6 | Primary sedimentation, oxidation ditch, secondary sedimentation, UV disinfection |
| W7 | Primary sedimentation, oxidation ditch, secondary sedimentation, UV disinfection |

**Table S2**: Concentration ranges (ng/L), median concentrations (ng/L), average concentrations (ng/L), and detection frequencies (%) of dissolved PPCPs in wastewater [1].

| **Analytes** | **Commercial use** | **Range** | **median** | **Average** | **Frequency** |
| --- | --- | --- | --- | --- | --- |
| 3-(4-methylbenzylidene)-camphor | UV filters | 1.22-47.0 | 10 | 12.4 | 100 |
| acetaminophen | NSAIDs | 640-5180 | 2580 | 2680 | 100 |
| acetophenone | Fragrance | BDL-96.2 | 23.4 | 28.1 | 71.4 |
| aspartame | Artificial sweetener | 0.488-304 | 17 | 28 | 100 |
| benzophenone-3 | UV filters | 1.36-12.9 | 6.2 | 6.47 | 100 |
| caffeine | Stimulant | 870-6220 | 2160 | 2470 | 100 |
| carbamazepine | Anticonvulsant | 2.30-40.6 | 11.3 | 12.2 | 100 |
| ciprofloxacin | Antibiotic | 6.58-84.8 | 30.6 | 33.2 | 100 |
| clenbuterol | β-sympathomimetic | 0.240-2.96 | 0.956 | 1.04 | 100 |
| clofibric acid | Lipid regulator | BDL-0.470 | BDL | 0.00959 | 2.04 |
| crotamiton | Antipruritic | 1.10-15.0 | 5.08 | 6.45 | 100 |
| cyclophosphamid | Antineoplastics | 0.107-4.64 | 0.486 | 1.04 | 100 |
| danofloxacin mesylate | Antibiotic | BDL-107 | 0.682 | 4.95 | 85.7 |
| diazepam | Anxiolytic | BDL-3.98 | 0.642 | 0.972 | 87.7 |
| enrofloxacin | Antibiotic | BDL-3.38 | 0.914 | 0.938 | 89.8 |
| ethenzamide | NSAIDs | BDL-2.16 | 0.332 | 0.374 | 83.7 |
| fenoprofen | NSAIDs | BDL-195 | 7.9 | 44.7 | 93.8 |
| fluoxetine | Antidepressant | BDL-61.8 | 19 | 20.6 | 97.9 |
| gemfibrozil | Lipid regulator | BDL-18.0 | 4.1 | 6.29 | 77.5 |
| ibuprofen | NSAIDs | 268-2240 | 628 | 811 | 100 |
| indomethacine | NSAIDs | 0.828-24.4 | 9.04 | 9.77 | 100 |
| ketoprofen | NSAIDs | 13.0-1030 | 236 | 299 | 100 |
| losartan | Hypertensive agent | BDL-4.12 | BDL | 0.539 | 34.6 |
| mefenamic acid | NSAIDs | 1.30-16.6 | 7.6 | 8.4 | 100 |
| methyl paraben | Preservative | BDL-154 | 45.8 | 46.1 | 91.8 |
| metoprolol | β-blockers | 3.32-328 | 51.4 | 90.6 | 100 |
| miconazole | Fungicide | 0.0848-7.82 | 1.56 | 1.97 | 100 |
| naproxen | NSAIDs | 1.63-20.4 | 11 | 11.4 | 100 |
| norfloxacin | Antibiotic | 123-2100 | 450 | 575 | 100 |
| ofloxacin | Antibiotic | 106-766 | 332 | 376 | 100 |
| oxytetracycline | Antibiotic | 39.6-398 | 152 | 161 | 100 |
| pirenzepine | Ulcer drug | BDL-1.45 | 0.104 | 0.149 | 93.9 |
| propranolol | β-blockers | 0.260-2.64 | 1.08 | 1.17 | 100 |
| propyl paraben | Preservative | 37.8-380 | 210 | 188 | 100 |
| propyphenazone | NSAIDs | 0.0850-7.70 | 0.55 | 1.45 | 100 |
| sarafloxacin hydrochloride | Antibiotic | BDL-1.61 | 0.105 | 0.151 | 46.9 |
| sildenafil | Sexual function agent | 0.254-8.84 | 2.92 | 3.41 | 100 |
| Sotalol | β-blockers | BDL-9.86 | 1.52 | 2.23 | 83.7 |
| tetracycline hydrochloride | Antibiotic | 23.0-197 | 68.8 | 78.6 | 100 |
| thiabendazole | Fungicide | BDL-0.798 | 0.252 | 0.263 | 75.5 |
| triclocarban | Antimicrobial | BDL-129 | 170 | 28.2 | 95.9 |
| triclosan | Antimicrobial | BDL-62.9 | BDL | 5.3 | 26.5 |

**Table S3**: PPCP mass load per capita of each classification (daily average) in influent [1].

|  | Influent (μg/d/inhabitant) | | | | | | |
| --- | --- | --- | --- | --- | --- | --- | --- |
| **Types** | W1 | W2 | W3 | W4 | W5 | W6 | W7 |
| Total PPCPs | 3086.83 | 2327.907 | 2530.332 | 1946.953 | 2222.245 | 1227.43 | 661.8642 |
| NSAIDs | 967.6492 | 806.4677 | 1050.049 | 620.7055 | 919.8385 | 414.7876 | 252.2573 |
| Antibiotics | 807.2898 | 899.5391 | 853.6979 | 826.2864 | 642.1482 | 485.5586 | 261.0711 |
| Stimulant | 1083.682 | 450.3999 | 498.3242 | 385.5678 | 539.2768 | 249.1304 | 109.1331 |
| Others PPCPs | 226.8697 | 170.8751 | 123.8332 | 120.8063 | 121.0568 | 78.8879 | 39.0966 |

**Figure**

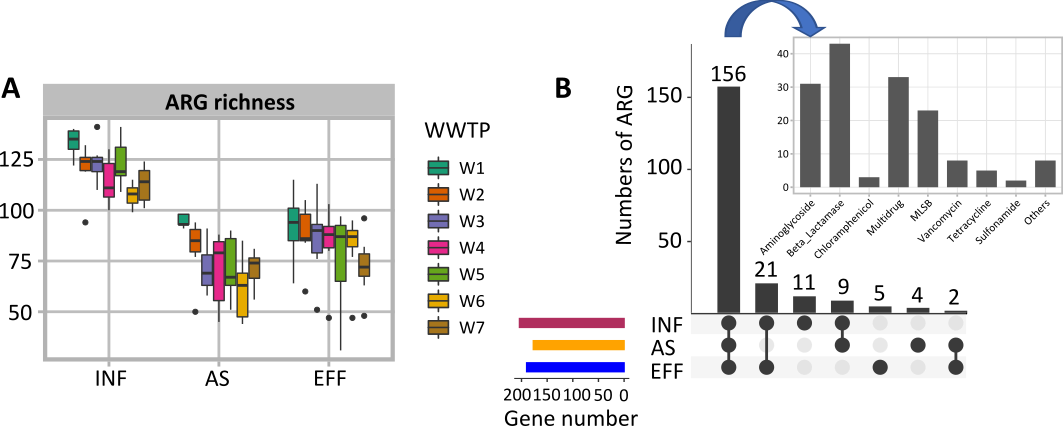

**Figure S1.** The number of ARGs detected in each treatment unit among seven WWTPs (A), unique and shared ARGs subtypes among different treatment units (B).

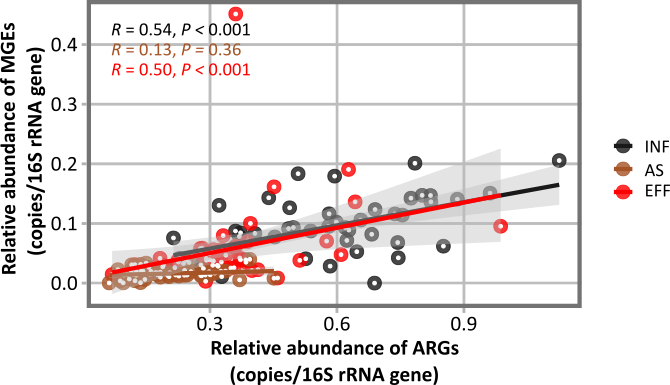

**Figure S2**. Spearman correlation between the relative abundance of ARGs and MGEs in WWTPs.

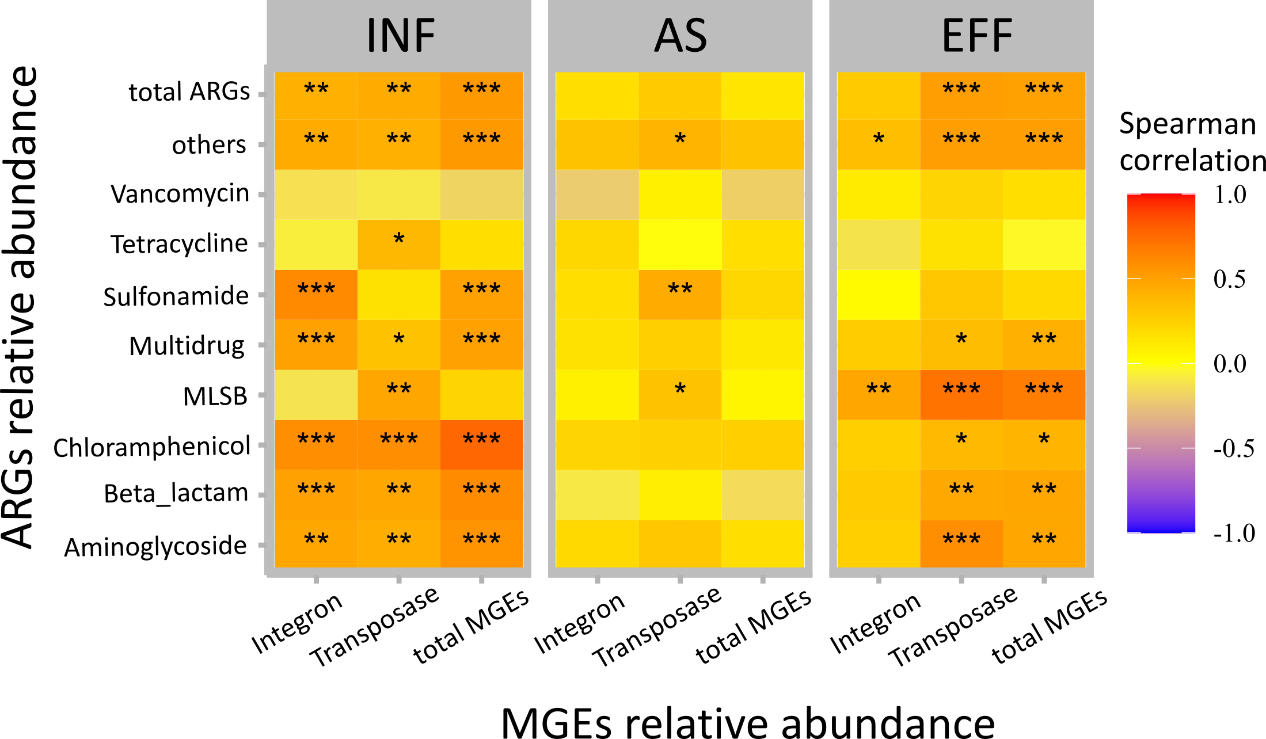

**Figure S3**. Spearman correlation analysis between the relative abundance of each ARG and MGE type in each treatment unit. Significance codes: *** *P* < 0.001, ** *P* < 0.01 and * *P* < 0.05.

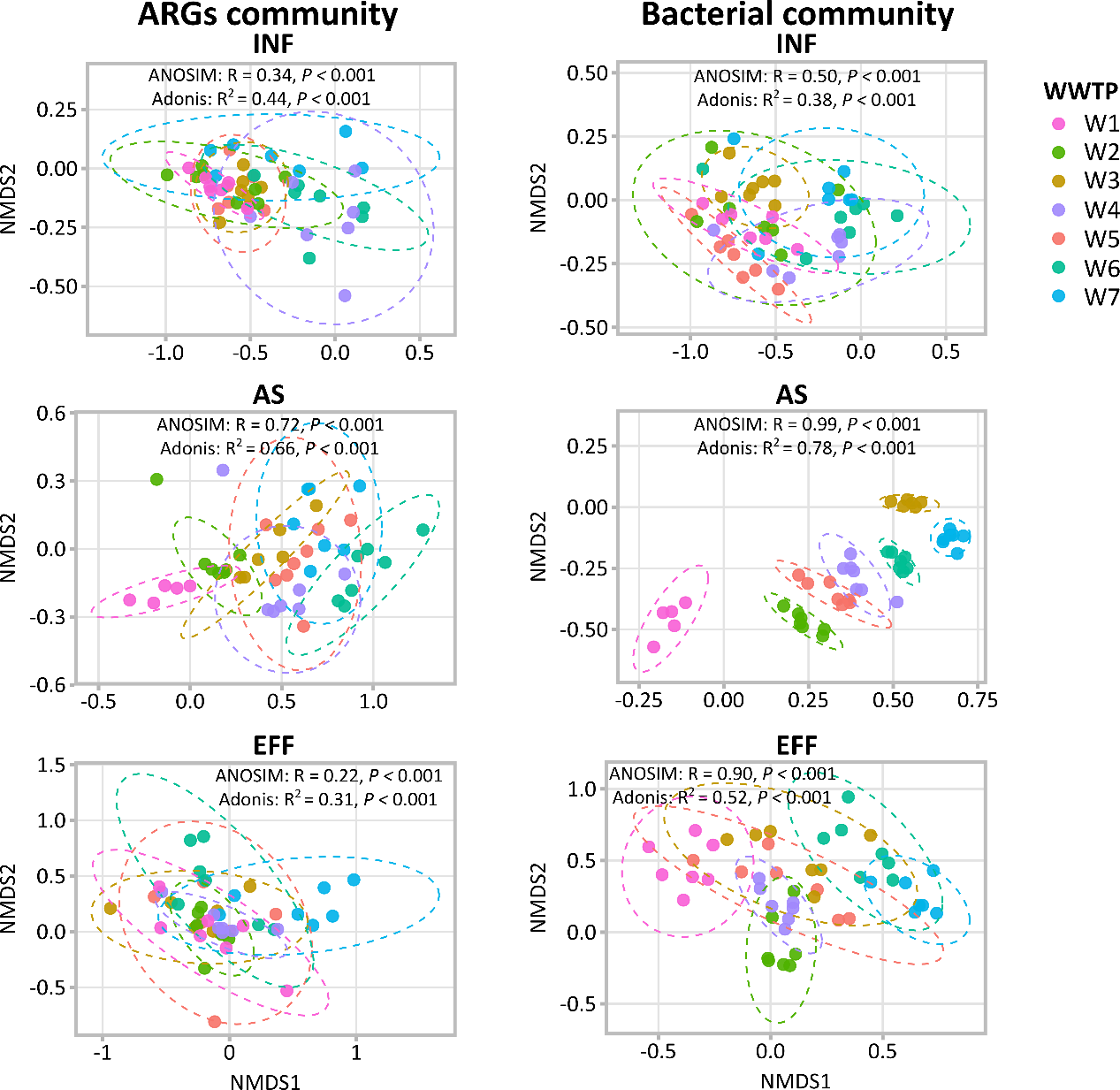

**Figure S4**. Non-metric multidimensional scaling (NMDS) ordination based on Bray-Curtis distance showed the distribution pattern of ARG and bacteria communities in each treatment unit. The level at 95% was drawn as an ellipse.

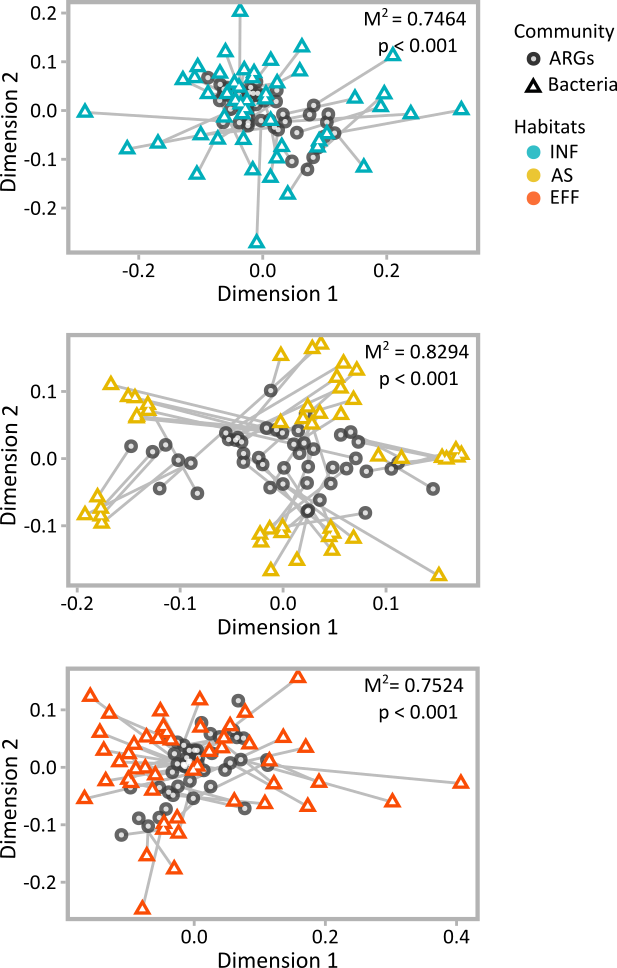

**Figure S5**. Procrustes analysis showing the significant correlation between the compositions of ARGs and bacterial communities from each treatment process.

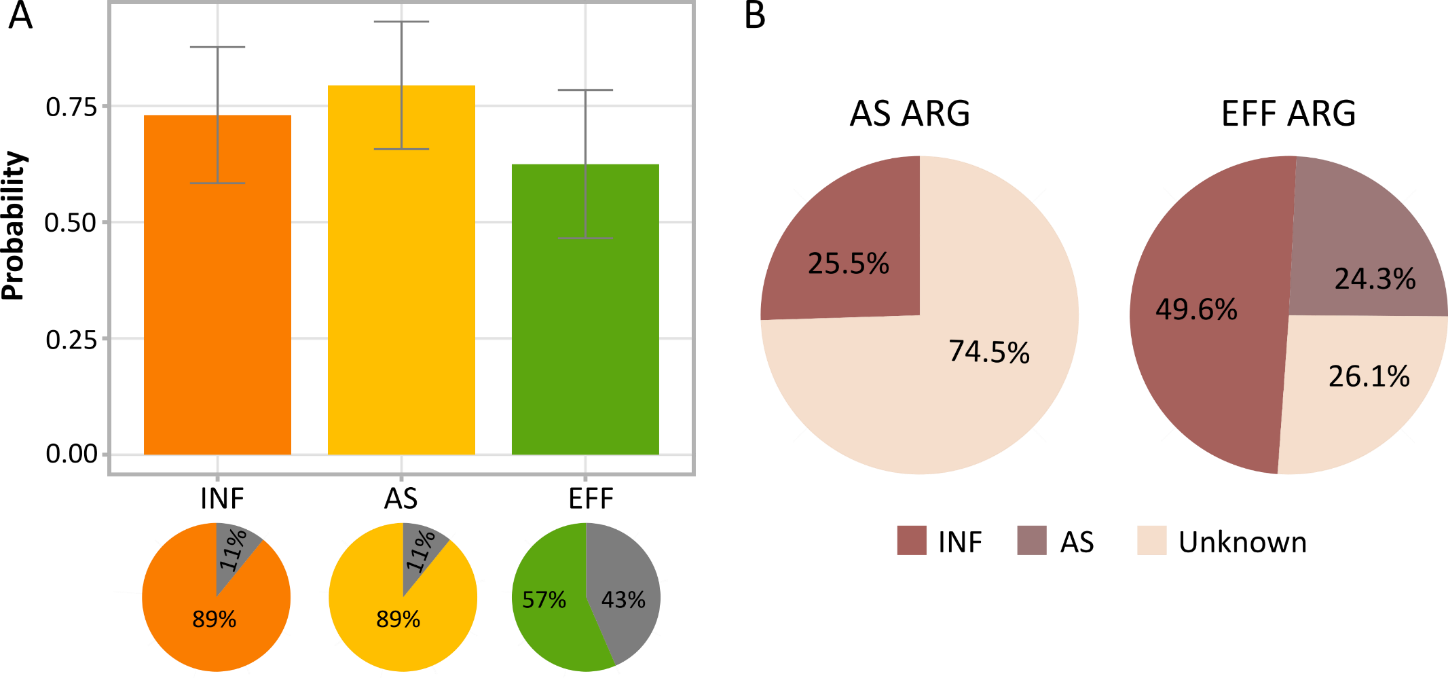

**Figure S6**. Ecotype prediction from ARGs community by SourceTracker using leave-one-out cross-validation (A), in which bar chart predicted probability and pie charts were correct prediction ratio of samples in the corresponding ecotype; (B) Results of SourceTracker in AS and EFF.

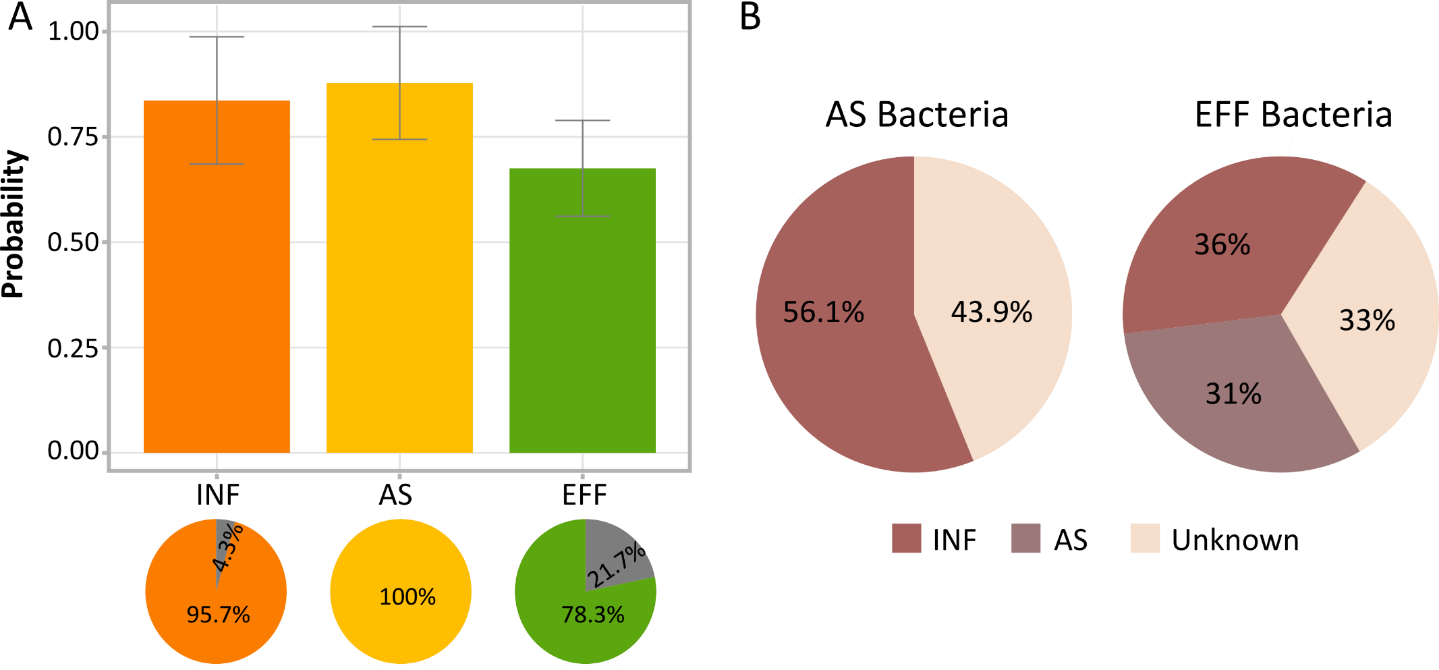

**Figure S7**. Ecotype prediction from bacteria community by SourceTracker using leave-one-out cross-validation (A), in which bar chart predicted probability and pie charts were correct prediction ratio of samples in the corresponding ecotype; (B) Results of SourceTracker in AS and EFF.

[1]	Y.Y. Wang, Y. Li, A. Hu, A. Rashid, M. Ashfaq, Y.Y. Wang, H. Wang, H. Luo, C.P. Yu, Q. Sun, Monitoring, mass balance and fate of pharmaceuticals and personal care products in seven wastewater treatment plants in Xiamen City, China, J. Hazard. Mater. 354 (2018) 81–90. https://doi.org/10.1016/j.jhazmat.2018.04.064.
